# Cerebellar mitochondrial dysfunction coincides with structural and behavioral abnormalities in 3q29Del mice

**DOI:** 10.64898/2026.09.25.754481

**Authors:** May K. Kretzer, Elizabeth Arnulfo Candelario, Ryan Montalvo, Xinyan Leng, Jonathan A. Coello, Zihui Ou, Aaron E. Pozo-Aranda, Jacob Ellegood, Jennifer G. Mulle, Zhen Yan, Ryan H. Purcell, Meike E. van der Heijden

## Abstract

3q29 deletion (3q29Del) is a genetic risk variant for autism spectrum disorder and schizophrenia that often results in developmental delays, cognitive disability, and impaired fine motor function. People with 3q29Del have reduced cerebellar volume, which correlates with symptom severity, and many 3q29Del-associated phenotypes also commonly occur after cerebellar injury or dysfunction. However, it is unknown whether the existing 3q29Del mouse model recapitulates the cerebellar dysfunction observed in humans. To characterize cerebellar phenotypes and uncover pathological differences in the 3q29Del mouse model, we investigated cerebellar structure, motor and vocal behaviors, protein expression, and mitochondrial function. We found uniformly reduced cerebellar volume in 3q29Del mice. Behavioral assays revealed vocal impairments in 3q29Del pups, fine motor impairments in adult mice, and reduced social mating calls in adult male mice. Proteomic analysis revealed enrichment of synaptic and mitochondrial proteins among the differentially expressed proteins in 3q29Del cerebellum tissue. Furthermore, mitochondria from 3q29Del mouse cerebellum displayed reduced oxygen flux and increased electron leak. These results recapitulate many human 3q29Del phenotypes in the 3q29Del mouse model and indicate mitochondrial dysfunction as a potential driver of 3q29Del pathology. Our findings also point to cerebellar involvement in 3q29Del phenotypes and provide a foundation for further research on cerebellar development in 3q29Del.

## 1. Introduction

Millions of people live with neurodevelopmental disorders such as autism spectrum disorder (ASD) and schizophrenia (SZ)^1^. Both ASD and SZ are highly heritable disorders, and to date hundreds of genetic variants have been identified that confer significant risk. However, there is no single causative variant^2^, and pathological effects of these risk variants on brain development and function are not fully understood. The hemizygous 3q29 deletion (3q29Del) is among the strongest known genetic risk factors for ASD and SZ^3,4^. 3q29Del is also associated with intellectual disability, executive function disorders, speech and motor delays, and graphomotor disability^3,5^, yet the precise underlying causes for these neurological disabilities remain unknown.

One potential mechanism implicated in ASD and SZ pathology is altered cellular metabolism, which could impair neuronal circuit formation and function particularly during energy-intensive phases of neurodevelopment. Mitochondrial dysfunction and disease are more prevalent among people with ASD^6,7^, and prior studies have observed mitochondrial dysfunction in 3q29Del as well as other ASD- and SZ-linked copy number variant conditions such as 22q11.2 deletion (22qDel)^8-13^.

Emerging evidence indicates that altered cerebellar development may also contribute to 3q29Del neuropsychiatric sequelae. Individuals with 3q29Del have reduced cerebellar volume, sometimes to the point of cerebellar hypoplasia, and the severity of the size reduction correlates with the severity of cognitive and motor symptoms^14,15^. Cerebellar circuits perform essential functions for motor control and speech^16,17^, both of which are impacted in 3q29Del^5,18^. These features are also broadly relevant as motor and speech abnormalities are often present in cases of ASD^19^ and SZ^20^. Conversely, cerebellar injury and dysfunction have been associated with increased prevalence for ASD^21-23^ and SZ^19,24,25^. Nevertheless, it remains largely unknown what cellular mechanisms link 3q29Del to altered cerebellar structure and impairments in cerebellum-associated behaviors.

The 3q29 region is conserved in mice, allowing the generation of a mouse model that recapitulates the genetic lesion and some behavioral phenotypes of 3q29Del^26^. While human research indicates cerebellar involvement in 3q29Del symptoms, the cerebellum is largely unexplored in the 3q29Del mouse model. To support mechanistic involvement of the cerebellum in 3q29Del, we set out to investigate whether cerebellar phenotypes are recapitulated in the mouse model. Our findings confirm the existence of cerebellar structural, molecular, and behavioral differences in the 3q29Del mouse model that closely resemble observations in people with 3q29Del.

## 2. Methods

### Animals

The relevant ethical guidelines for animal testing were followed at all times, and all husbandry, experimental, and euthanasia protocols were approved by the Institutional Animal Care and Use Committee (IACUC). 3q29Del mice were provided by the Emory University Mouse Transgenic and Gene Targeting Core Facility (RRID:SCR_023535) and bred in-house on a C57BL/6N background (Charles River). Genotypes of offspring were confirmed with PCR of tail or ear clippings. Male and female mice were studied in all experiments except the courtship assay.

### Tissue fixation and immunostaining

Perfusion and tissue fixation were performed by thoroughly anesthetizing mice with isoflurane via inhalation and perfusing the whole body with 1X phosphate-buffered saline (PBS), then with 4% paraformaldehyde (PFA) diluted in PBS. The whole brain was then dissected and stored in 4% PFA overnight, then placed in increasing concentrations of sucrose in 24-hour time durations: 10% sucrose, then 20%, then 30%. After 24 hours of storage in 30% sucrose, brains were embedded in OCT compound and frozen at -80C. Brains were sectioned sagittally at 40um on a cryostat, then stored in 0.1% sodium azide at 4C.

For immunohistochemistry, free-floating sections were incubated in a solution of 10% normal goat serum (NGS) diluted in PBS-T for 1 hour at room temperature, then incubated in the same solution containing primary antibodies for 12-18 hours at room temperature. Sections were then washed with PBS-T 3 times and incubated in PBS-T containing secondary antibodies for 2 hours at room temperature. Sections were then washed with PBS-T 3 times again and mounted onto slides, using VectaShield mounting medium. Negative controls underwent these exact same steps except that there was no primary antibody in the 10% NGS solution during the primary incubation period.

### Imaging and image analysis

Images of whole-cerebellum sections were captured using a Leica fluorescent microscope at 20X magnification. Surface area of sections were calculated using the polygon tool in ImageJ to create a detailed outline of the section and calculate the area within. Sections were classified as vermis, paravermis, or hemisphere by agreement of at least two researchers, and sections were excluded from analysis if they were too damaged or folded to calculate area, or if they were from an extremely lateral part of the cerebellum. All researchers performing quantifications were blinded to the genotypes of the mice each section belonged to. Images were corrected for brightness and contrast using ImageJ for ease of properly capturing the section outline when applicable, and for use in figures.

### Magnetic Resonance Imaging

A multi-channel 7.0 Tesla MRI scanner (Agilent Inc., Palo Alto, CA) was used to image the brains within their skulls. Sixteen custom-built solenoid coils were used to image the brains in parallel^27,28^.

#### Anatomical scan

In order to detect volumetric changes, we used the following parameters for the MRI scan: T2 - weighted, 3-D fast spin-echo sequence, with a cylindrical acquisition of k-space, a TR of 350 ms, and TEs of 12 ms per echo for 6 echoes, field-of-view equaled to 20 × 20 × 25 mm3 and matrix size equaled to 504 × 504 × 630. Our parameters output an image with 0.040 mm isotropic voxels. The total imaging time was 14 hours^29^.

#### MRI registration and analysis

To visualize and compare any changes in the mouse brains the images are linearly (6 followed by 12 parameter) and non-linearly registered together. Registrations were preformed with a combination of mni_autoreg tools^30^. and ANTS (advanced normalization tools)^31,32^. All scans are then resampled with the appropriate transform and averaged to create a population atlas representing the average anatomy of the study sample. The result of the registration is to have all images deformed into alignment with each other in an unbiased fashion. For the volume measurements, this allows for the analysis of the deformations needed to take each individual mouse’s anatomy into this final atlas space, the goal being to model how the deformation fields relate to genotype^33,34^. The Jacobian determinants of the deformation fields are then calculated as measures of volume at each voxel. Significant volume differences can then be calculated by warping a pre-existing classified MRI atlas onto the population atlas, which allows for the volume of 182 different segmented structures encompassing cortical lobes, large white matter structures (i.e. corpus callosum), ventricles, cerebellum, brain stem, and olfactory bulbs^35-38^ to be assessed in all brains. Further, these measurements can be examined on a voxel-wise basis in order to localize the differences found within regions or across the brain. Multiple comparisons in this study were controlled for using the False Discovery Rate^39^.

### Behavioral assays

#### Surface righting reflex

To assess surface righting reflexes, the researcher turned mouse pups over onto their backs and timed latency to fully reorient themselves to standing on their paws; time was recorded to the nearest tenth of a second^40^.

#### Negative geotaxis reflex

To assess negative geotaxis reflexes, the researcher placed mouse pups onto a ∼40 degree incline facing downwards and timed how long it took them to turn around. The timer was stopped once the animal’s head was oriented at an angle greater than 90 degrees (past perpendicular) to where they were initially facing, and time was recorded to the nearest tenth of a second^40^.

#### Pup vocalization recordings

To assess vocalizations, mouse pups were taken from their home cage and immediately placed in a soundproof chamber with a microphone^40^. Vocalizations were recorded for two minutes, and the number and duration of these vocalizations were reported by SonoTrack software. Recordings were then computationally classified and analyzed (SonoTrack). Vocalizations were classified into 14 different types: short, flat, up, down, chevron, U-shape, trailing, step down, step up, step double, complex-3, complex-4, complex-5, and complex-5+. There were two additional categories for undefined vocalizations—undefined long and undefined short—for a total of 16 possible classifications. All vocalizations not classified as “short” were considered complex.

#### Courtship vocalization assay

Adult (10-12 week) male mice were habituated to the USV chamber alone in a clean cage with no bedding for 10 minutes prior to testing, then for a 2 minute period immediately before testing. Mice were then placed in the chamber with a novel female mouse of a similar age for 2 minutes. This was repeated for 2 trials, and the trial in which the mouse vocalized the most was used for analysis (trial 1 was used in ties). Vocalizations were analyzed using the same method as described in pup vocalizations. Only male mice were used because male mice vocalize to novel females, but female mice do not vocalize to novel males^41,42^.

#### Rotarod

Adult (6-12 week) mice were placed on a surface rotating initially at 5 rotations per minute (rpm) that gradually accelerated to 40rpm over the course of 300 seconds^43^. The time each animal remained on the surface before failing the trial was recorded. A mouse was considered to have failed the trial if it fell from the rotarod or clung to it for two full rotations without returning to a walking pace on top of the surface. Three trials were conducted each day over the course of three consecutive days, with a short rest period in between trials. Data are reported as the average of three trials on each day.

#### String pull

Adult (7-10 week) mice were food-restricted (2.5g chow for males, 2g for females) for three days prior to testing. To habituate the mice to strings and the food reward, they were placed in a cage containing the hanging ends of strings with a reward (fruit loop) tied to the other end one day prior to testing. There were ten strings 20-80cm long in each habituation cage, and metal bars at the top prevented mice from pulling the food fully into the cages. Tests were performed in a clear container with the reward at the end of a 100cm string^44^, and videos were recorded for three trials per mouse. If the animal did not pull the string within twenty minutes, it was considered a failed trial. Video recordings were manually scored by blinded researchers to determine error rate. Possible errors are a *partial miss*, pulling twice with the same paw, a *double pull*, pulling with both paws at once, and a *full miss*, reaching for the string and failing to grasp it.

### Proteomics

Brain samples were homogenized in 8M Urea lysis buffer and water bath sonicated for 15 mins. BCA was performed and 50ug protein used for digestion. Samples were reduced in 5 mM DTT at ambient temperature for 30 min and then alkylated with 10mM IAA in the dark for 30 min. The samples were then digested with 1ug of Lysyl endopeptidase (Wako) and 2ug trypsin (Thermo) overnight at room temperature. The peptide solutions were then acidified to 1% FA and cleaned with microelute HLB columns. HLB columns were conditioned with 500uL of methanol and equilibrated twice with 500uL of 1% formic acid. Samples were loaded onto the HLB column and then washed twice with 500ul of 1% formic acid. Finally, the samples were eluted with 100uL of 50% acetonitrile. The eluates were then dried to completeness using a SpeedVac (LabConco).

Each sample was resuspended in 50ul of loading buffer (0.1% FA) and 1ul was analyzed by liquid chromatography coupled to tandem mass spectrometry. Peptide eluents were separated on Water’s CSH fused silica column (8 cm × 150 μM internal diameter packed with CSH 1.7um resin) by a Vanquish Neo (Thermo). Buffer A was water with 0.1% (vol/vol) formic acid, and buffer B was 80% (vol/vol) acetonitrile in water with 0.1% (vol/vol) formic acid. Elution was performed over a 17 min gradient, up to 50% solvent B. Peptides were monitored on a Orbitrap Astral mass spectrometer (Thermo) fitted with a high-field asymmetric waveform ion mobility spectrometry (FAIMS Pro) ion mobility source (Thermo). One compensation voltage of -35 was chosen for the FAIMS. Each cycle consisted of one full scan (MS1) performed with an m/z range of 380-980 at 240,000 resolution, 500% AGC and 3 ms injection time. The higher energy collision-induced dissociation (HCD) DIA scans were collected with 2 m/z isolation windows over the entire precursor range (380-980 m/z) with a time of 0.6 seconds and 2.5 ms injection time. Collision energy was set to 27% and scan range set to 150 - 2000 m/z. Spectronaut (version 19.7.250203) was used to search all raw files in default library free mode using a mouse uniprot database (downloaded 032024 and supplemented with custom mutant sequences – total entries 17204). All parameters were kept at default.

### Evaluation of Mitochondrial Respiration and Electron Leak

#### Mitochondria isolation

Mitochondria were isolated from fresh cerebellum utilizing previously confirmed methods^45^. In brief, fresh samples were placed immediately in ice cold mitochondria isolation buffer [MIB: BSA (2 mg/mL), Sucrose (70 mM), Mannitol (210 mM), HEPES (5 mM), EGTA (1 mM), pH 7.1] and briefly processed with a saw-tooth homogenizer over an ice bath. Homogenate underwent two rounds of differential centrifugation at 4ºC at 800xg and 9000xg for 10 minutes each to isolate a mitochondrial pellet and cytosolic fraction. Mitochondria were resuspended in MIB without BSA and quantified utilizing the Bradford method to allow for normalization of protein load for evaluation of *J*O_2_ and *J*H_2_O_2_.

##### Mitochondrial respiratory flux (*J*O_2_) and reactive oxygen species flux (*J*H_2_O_2_)

were evaluated underneath the creatine kinase clamp^45,46^. Contrary to traditional methods, the creatine kinase clamp utilizes the creatine kinase enzyme to maintain stoichiometric ratios of ATP:ADP and PCr:Cr within physiological relationships that reflect *in vivo* conditions. Measures of *J*O_2_ were performed in an Oroboros O2k (Innsbruck, Austria) maintained at 37ºC with a magnetic stir set at 500 rpm in 0.5mL chambers. Chambers were calibrated with buffer D (BxD) [KMES (105 mM), KCl (30 mM), EGTA (1 mM), KH2PO4 (10 mM), MgCl2-6H2O (5 mM), 0.05% BSA, pH 7.1, solubility factor 0.966] to assess R1 (air saturation, ∼200 µM O2) and R0 (zero oxygen) with sodium hydrosulfide. Respiratory assessments of oxygen flux (JO2 (pmol/sec/mg)) were made with BxD supplemented with 5mM creatine monohydrate and 15μg of mitochondrial cerebellar protein, normalized as described above.

Mitochondrial integrity was confirmed with addition of cytochrome c (0.005mM) with an *a priori* threshold of 15%, which did not require the exclusions of any samples from this data set. Malate (2.5mM), pyruvate (5mM), and glutamate (5mM) were utilized for evaluation of non-phosphorylating respiration and as substrates for maximal respiration at ΔG_ATP_ -12.94 as calculated using a freely available github calculator (https://dmpio.github.io/bioenergetic-calculators/ck_clamp/) with conditions of 37ºC temperature, 170mM ionic strength, 5mM creatine, 10mM phosphate, and pH 7.1 based on parameters referenced here (https://github.com/dmpio/bioenergetic-calculators/blob/master/jupyter_notebook/creatine-kinase-clamp.ipynb). *J*H_2_O_2_ was evaluated in a fluorolog-QM (Horiba Scientific, Edison, NJ) outfitted with a 4-rotor turret system (Quantum Northwest, Liberty Lake, WA) using the Amplex Ultra Red (AUR) Horseradish Peroxidase (HRP) assay; AUR is converted to H_2_O_2_ after reacting with HRP for fluorometric detection. This reaction occurs in the same buffer D supplemented with Cr as well as superoxide dismutase (SOD) to ensure complete conversion of free radicals (•O_2_) to hydrogen peroxide (H_2_O_2_) and auranofin to ensure inhibition of ROS scavenging^46^. Rates of *J*H_2_O_2_ were fit to a standard curve and electron leak was calculated by dividing the oxygen consumed by the AUR HRP reaction by the oxygen consumed by OXPHOS and is expressed as a percentage. Standard student T tests were performed between groups to identify differences with significance set at p<0.05 vs WT littermate controls.

### Statistical Analyses

All statistical analyses were performed with GraphPad Prism software, and data are all presented as means. Behavioral data were analyzed with two-tailed student t-tests or Mann-Whitney tests when distributions were not normal, and cerebellar size data were analyzed with nested t-tests. For all statistical tests, results were considered statistically significant if p < 0.05.

## 3. Results

### 3.1: Reduced cerebellar size in 3q29Del mice

Reduced cerebellar volume is a common neuroanatomical observation in people with 3q29Del^14,15^, but a previous study did not observe any gross cerebellar abnormalities in mice^26^. To investigate whether the 3q29Del mouse model recapitulates reduced cerebellar volume, we measured cerebellar volume of 3q29Del and wild-type littermate control mice. Structural MRI of the entire cerebellum and cerebellar subregions in adult mice revealed significant volumetric reductions (Fig. 1a). This effect was consistent across the cerebellar cortex, hemispheric regions, and vermal regions (Fig. 1b-d), indicating that 3q29Del causes a global cerebellar size reduction. This is largely consistent with cerebellar phenotypes seen in human study participants with 3q29Del: most subregions of the cerebellum are significantly reduced in size, although there is variability between subregions and individuals^14,15^.

**Figure 1:**
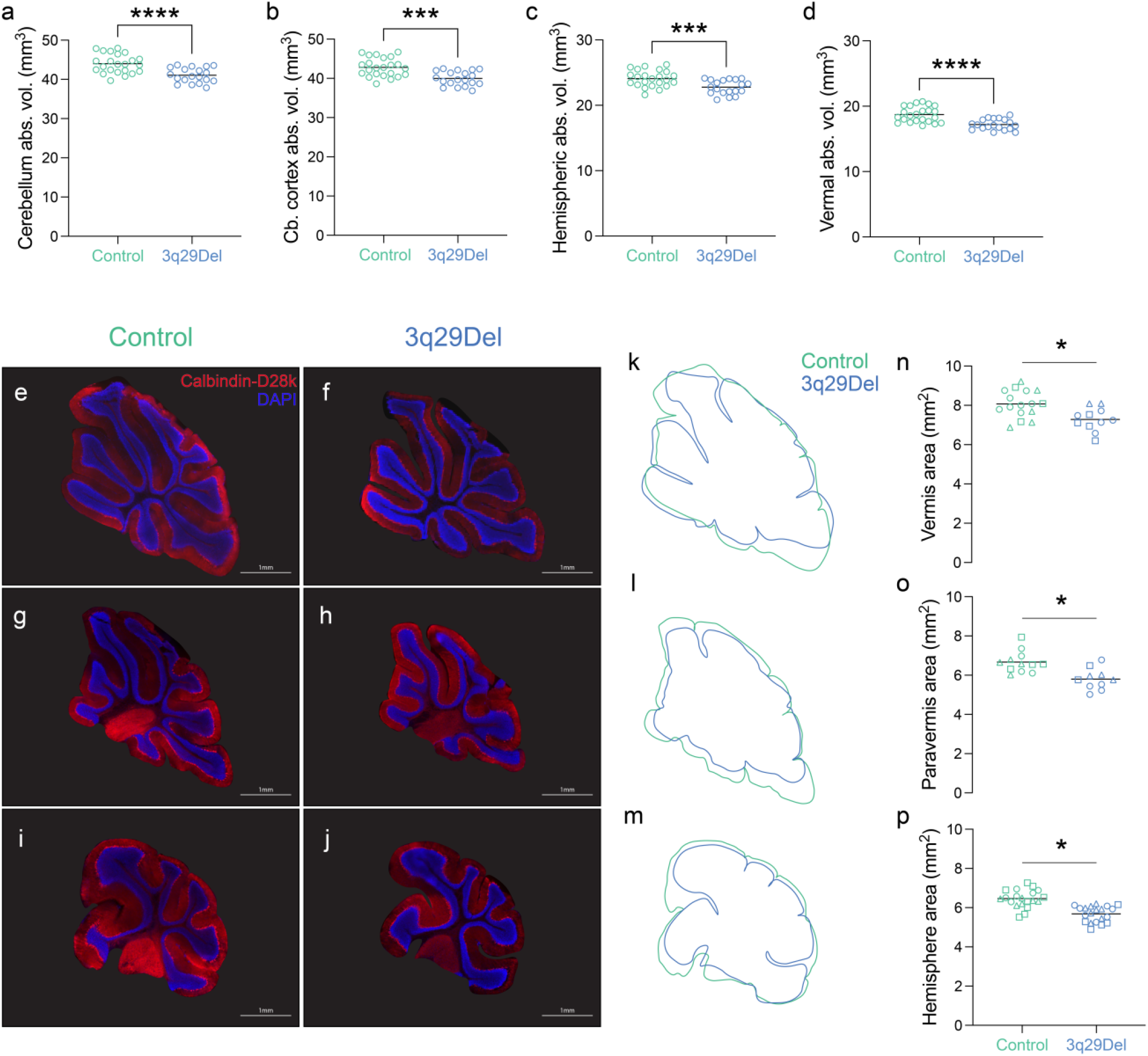
Cerebellar size reduction in 3q29Del mice. Absolute volume as measured by structural MRI of the cerebellum (a), cerebellar cortex (b), hemispheric regions (c), and vermal regions (d). Representative images of vermis (e-f), paravermis (g-h), and hemisphere (i-j) sections stained with DAPI and calbindin to indicate Purkinje cells and distinguish layers. Outline overlays of these images (k-m) and sectional surface area (n-p). N=3 mice per group, 3-5 sections per mouse; each data point represents one section and each shape represents one mouse. Nested t-test results-Vermis: t(4)=3.1 p=0.036. Paravermis: t(4)=3.7, p=0.021. Hemisphere: t(4)=4.7p=0.0095.

Medial (vermal) regions develop slightly earlier than more lateral (paravermal and hemispheric) regions, and projections from the cerebellar cortex to the deep cerebellar nuclei are organized in a mediolateral manner^47,48^. Thus, it is important to determine whether any mediolateral regions are differentially affected by 3q29Del. To investigate regional differences in finer detail, we approximated volume with the surface area of sagittal cross-sections of P28 mouse cerebella. We categorized each section by mediolateral region: vermis (most medial), paravermis (slightly lateral), and hemisphere (most lateral). These measurements also revealed a significant size reduction of approximately 10% in the cross-sectional surface area of each mediolateral region of the cerebellum (Fig. 1e-p), a similar effect size as we observed with the structural MRI. Also in agreement with the MRI results, the size difference appears to be uniform, indicating that 3q29Del has a consistent effect on cerebellar growth across the mediolateral axis.

The volumetric and surface area quantifications were conducted in separate laboratories in distinct mouse colonies, supporting the reproducibility of this result. These findings provide strong evidence that this 3q29Del mouse model recapitulates the human phenotype of reduced cerebellar volume, indicating cerebellar deficits are a cross-species impact of the 3q29 deletion. These data support the utility of the 3q29 deletion mouse model as a resource for studying mechanisms by which 3q29del cerebellar changes contribute to risk for neurodevelopmental phenotypes.

### 3.2: Reduced vocalization complexity in 3q29Del mouse pups

Next, we set out to investigate whether the reduced cerebellar size may contribute to cerebellum and 3q29Del associated behavioral deficits. Children with 3q29Del often exhibit delays in motor function and speech^5^, which can also occur in infants with perinatal cerebellar injury^21^. Previous studies have tested motor coordination in adult 3q29Del mice and found no impairment^26,49^, but did not test developing pups. A recent study indicated that 3q29Del pups vocalize less in isolation^50^. Yet, it is unknown whether 3q29Del also alters bioacoustic parameters that are sensitive to cerebellar circuit function^51^, like vocalization length, tonality, loudness, or complexity. We therefore investigated whether 3q29Del pups showed abnormal early postnatal motor and vocal behaviors.

First, we tested two motor reflexes: surface righting reflex and negative geotaxis reflex (Fig. 2a,c), as prior research has found that cerebellum-specific circuit manipulations impair these reflexes^40,43^. We tested the mice at postnatal day 7 (P7), at which point these reflexes exist, but motor circuits are still developing and may impede response time in cases of developmental delay.

**Figure 2:**
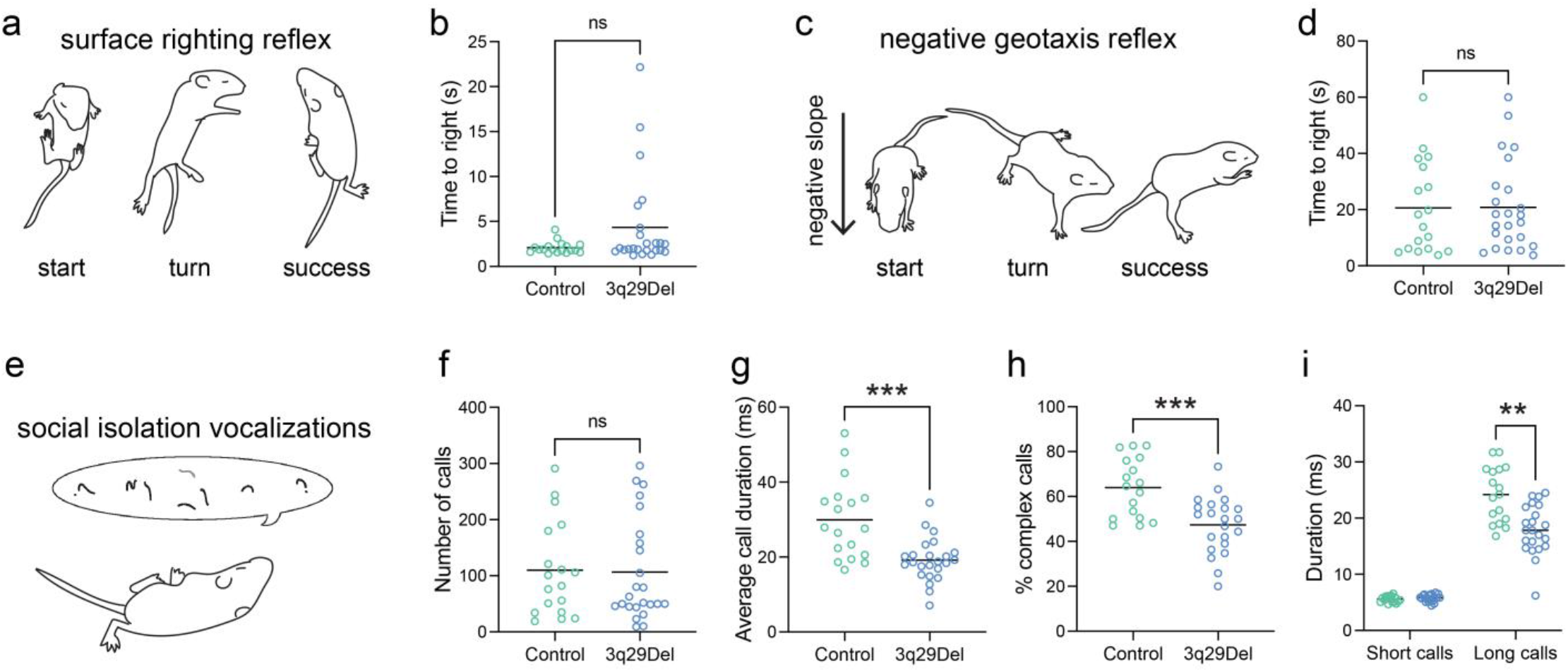
At P7, 3q29Del mouse pups exhibit behavioral differences. Illustration of righting reflex assay (a), time to right (b); Mann-Whitney test: U=163, p=0.18. Illustration of negative geotaxis assay (c), time to turn (d); unpaired t test: t(40)=0.032, p=0.97. Illustration of vocalization assay (e). Average number of calls (f); unpaired t test: t(40)=0.12, p=0.91. Average call duration (g); unpaired t test: t(40)=4.2 p = 0.0008. Percentage of complex vocalizations (h); unpaired t test: t(40)=4.0, p=0.0003. Average durations of short (t(40)=1.0, p=0.30) and long (t(37)=4.2, p=0.0002) vocalizations (i). Control: N = 18 (M=9, F=9), 3q29Del: N=24 (M=9, F=15).

A few 3q29Del pups took an abnormally long time to right themselves (Fig. 2b), but most were close to the control mean in response time. Overall, 3q29Del pups were not significantly different from control animals in either reflex. This suggests that 3q29Del does not critically impair early postnatal motor milestones in 3q29Del mice.

Next, we studied pup vocalizations in the social isolation paradigm. Although this paradigm is often used to test inclination to vocalize (number of vocalizations), we were particularly interested in the qualitative measures and power-spectrums of the vocalizations given the speech delay observed in children with 3q29Del and prior findings that cerebellar manipulations can cause altered vocalizations in mice^40,43,51,52^. We recorded and analyzed vocalizations from P7 mouse pups with SonoTrack software, which counted and categorized calls based on spectrotemporal features such as shape, syllables, and pitch changes. We found that 3q29Del and control pups produced similar numbers of vocalizations (Fig. 2f), which reflects that these 3q29Del pups are both able and inclined to vocalize when separated from their mother. However, we noted that the average call duration of 3q29Del pups’ calls was shorter than that of controls (Fig. 2g). To determine whether this reduction in call duration was affected by call type (i.e. more short calls and fewer long), we compared the categorization of each vocalization. Among the many distinct types of mouse vocalizations, the simplest and most common is the short call. We grouped the pups’ vocalizations into short calls and all other call types (referred to here as “complex calls”) and found that 3q29Del pups make a lower percentage of complex calls (Fig. 2h).

3q29Del pups make fewer complex calls, but are the complex calls they produce similar to controls? To test this, we grouped calls by duration (short:<20ms, and complex: ≥20ms) and quantified the average duration of each group. We found no difference in length of the shorter calls, but the longer calls made by 3q29Del pups were still significantly shorter on average (Fig. 2i). This result indicates that in addition to making fewer complex calls, the complex calls that 3q29Del pups produce are shorter. Our results suggest that 3q29Del pups do not demonstrate disability or inclination to vocalize, but may have difficulty making more complex sounds, which could reflect impaired vocal-motor coordination.

### 3.3: Deficits in fine motor coordination and social vocalizations in adult 3q29Del mice

Some individuals with 3q29Del display motor impairments^15^, which may be caused by reduced cerebellar size. Several studies report high prevalence of fine motor deficits, such as writing disabilities, in the 3q29Del population^5,14,18^, and some people with 3q29Del exhibit gait ataxia^53^. To thoroughly assess motor capabilities in adult 3q29Del mice, we performed behavioral assays designed to test gross and fine motor skills.

To test balance, gross motor coordination, and motor learning, we administered a rotarod test (Fig. 3a). Our lab and others have shown that performance on this test is disrupted by cerebellar manipulations ^40,43,54,55^, and various mouse models of ASD have also been observed to perform poorly on this test^56-58^. 3q29Del mice, however, performed similarly to controls, indicating no severe deficits in balance or motor learning.

**Figure 3:**
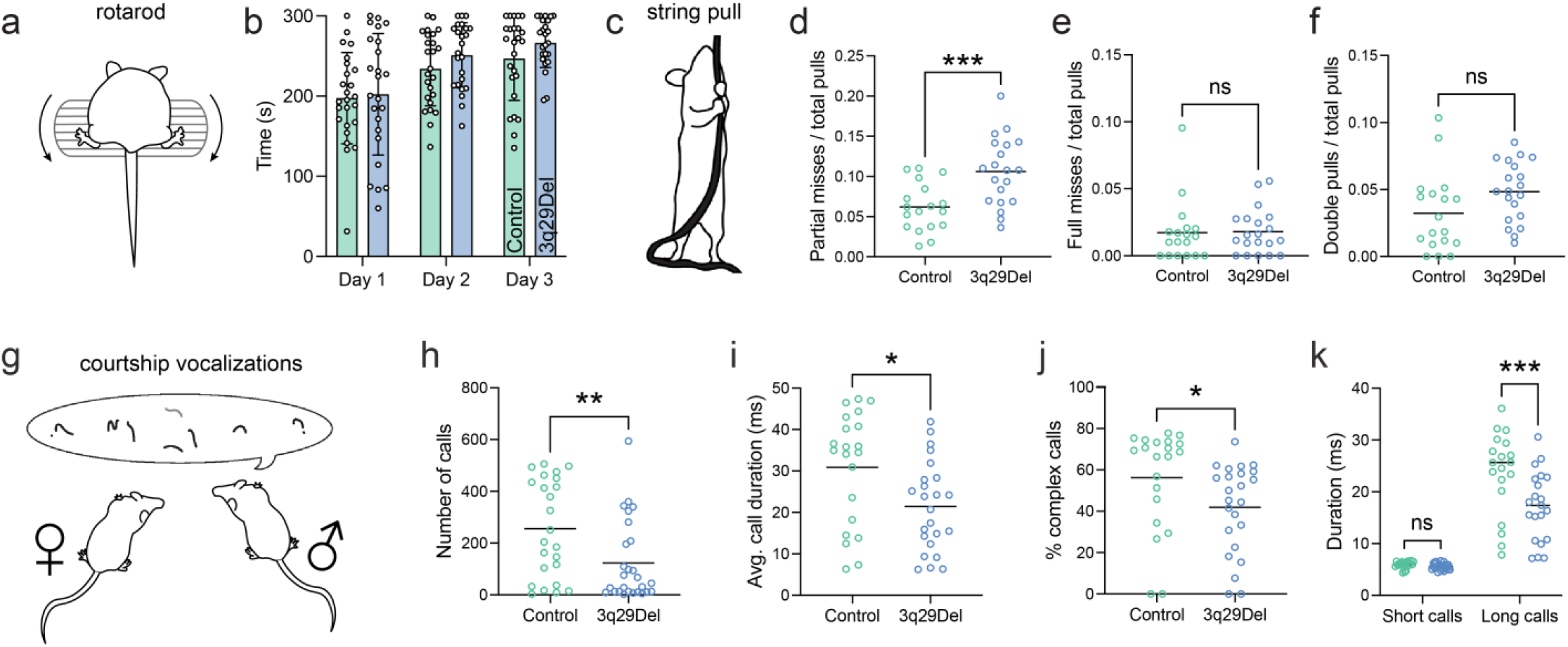
Adult 3q29Del mice have impaired fine motor coordination and social vocalizations. Illustration of rotarod assay (a), time to rotarod failure (b). Control: N=25 (M=11, F=14), 3q29Del: N=24 (M=13, F=11). Illustration of string pull assay (c), partial miss rate (d); unpaired t test: t(36)=3.7, p=0.0008. Full miss rate (e); t(36)=0.12, p=0.91. Double pull rate (f); t(36)=1.9, p=0.064. Controls: N=18 (M=10, F=8), 3q29Del: N=20 (M=12, F=8). Illustration of courtship vocalization assay (g), number of calls (h); Mann-Whitney test: U=179.5, p=0.0057. Average call duration (i); unpaired t test: t(43)=2.6, p=0.012. Percentage of complex calls (j); unpaired t test: t(43)=2.1, p=0.042. Average durations of short (t(43)=1.8, p=0.075) and long (t(38)=2.8, p=0.0076) calls (k). Control: N=21, 3q29Del: N=24 (all male).

To test whether 3q29Del mice have impaired fine motor coordination, a much more common motor phenotype in people with 3q29Del, we administered a string pull test (Fig. 3c). In this test, mice pull down a string with a food reward at the end. Recordings of this test were manually scored for mistakes. As efficient string pulling is done paw-over-paw, deviations from that pattern were counted as mistakes^59^. (Supplemental video 1). Partial miss errors, in which a mouse pulls with the same paw twice in a row, were made at a significantly higher rate in 3q29Del animals than controls (Fig. 3d). This finding indicates a significant fine motor impairment in 3q29Del mice. The increased partial miss rate, but lack of significant differences in other error types, suggests that 3q29Del mice are less skilled at coordinating small movements with both front paws, but not so uncoordinated that they use both paws to pull, as has been observed in mice with atypical cerebellar circuitry^54^. Because we observed vocal deficiencies in 3q29Del pups, we tested whether these impairments persisted into adulthood. Measuring vocalizations in adult mice is more difficult than in pups: adult mice vocalize very little, if at all, in isolation, and most social assays involve multiple possibly-vocalizing mice that could not be differentiated with our equipment. Thus, we elected to use a courtship assay, as males vocalize during courtship and females do not. This enabled accurate recording and attribution of calls but limited the data to only male mice in a courtship scenario. Use of this assay also introduced a significant social component to be taken into consideration. We found that in a courtship setting, male 3q29Del mice vocalized significantly less often than controls (Fig. 3h). Additionally, and consistent with our findings in pups, 3q29Del mice produced a significantly lower percentage of complex calls, and their longer calls were shorter than controls’ (Fig. 3j-k). These data could indicate that the potential vocal disabilities observed in 3q29Del pups persist into adulthood. Our behavioral data reflect social, vocal, and fine motor deficits in 3q29Del mice that may be the result of cerebellar dysfunction.

### 3.4: Mitochondrial dysfunction in 3q29Del cerebellum

To investigate potential molecular drivers of altered cerebellar structure in 3q29Del, we analyzed the proteomes of bulk cerebellar tissue in adult mice (N=9/genotype, age 6-11 weeks, all male). There was no significant difference in age between groups (Control=56.5 days, 3q29Del=57.1 days, p=0.76). We quantified the relative expression of 8,442 proteins. Proteins with absolute log2 fold change greater than 0.1 and FDR-corrected q-value less than 0.05 were considered differentially abundant between genotypes. 1,457 proteins met these criteria (Fig. 4b, 790 down). 10 of 22 proteins of the homologous 3q29Del interval were detected and quantified (remaining proteins were not consistently detected across all samples). Each of these ten proteins were found to be significantly decreased, most by the expected ∼50% reduction to match gene copy number (Fig. 4c), indicating that 3q29Del consistently changed protein abundance of encoded genes in cerebellum. Pathway analysis of all differentially abundant proteins that met a strict statistical cut-off (q<0.001) revealed significant enrichment for synaptic and mitochondrial ontologies including GO:0045202 *synapse*, GO:0098978 *glutamatergic synapse*, GO:0005739 *mitochondrion*, and GO:0006119 *oxidative phosphorylation* (Fig. 4d). We further investigated synaptic enrichment by performing SynGO analysis (Fig. 4e-f) and indeed found significant enrichment for both pre- and post-synaptic components with some additional evidence for vesicle-related processes.

**Figure 4:**
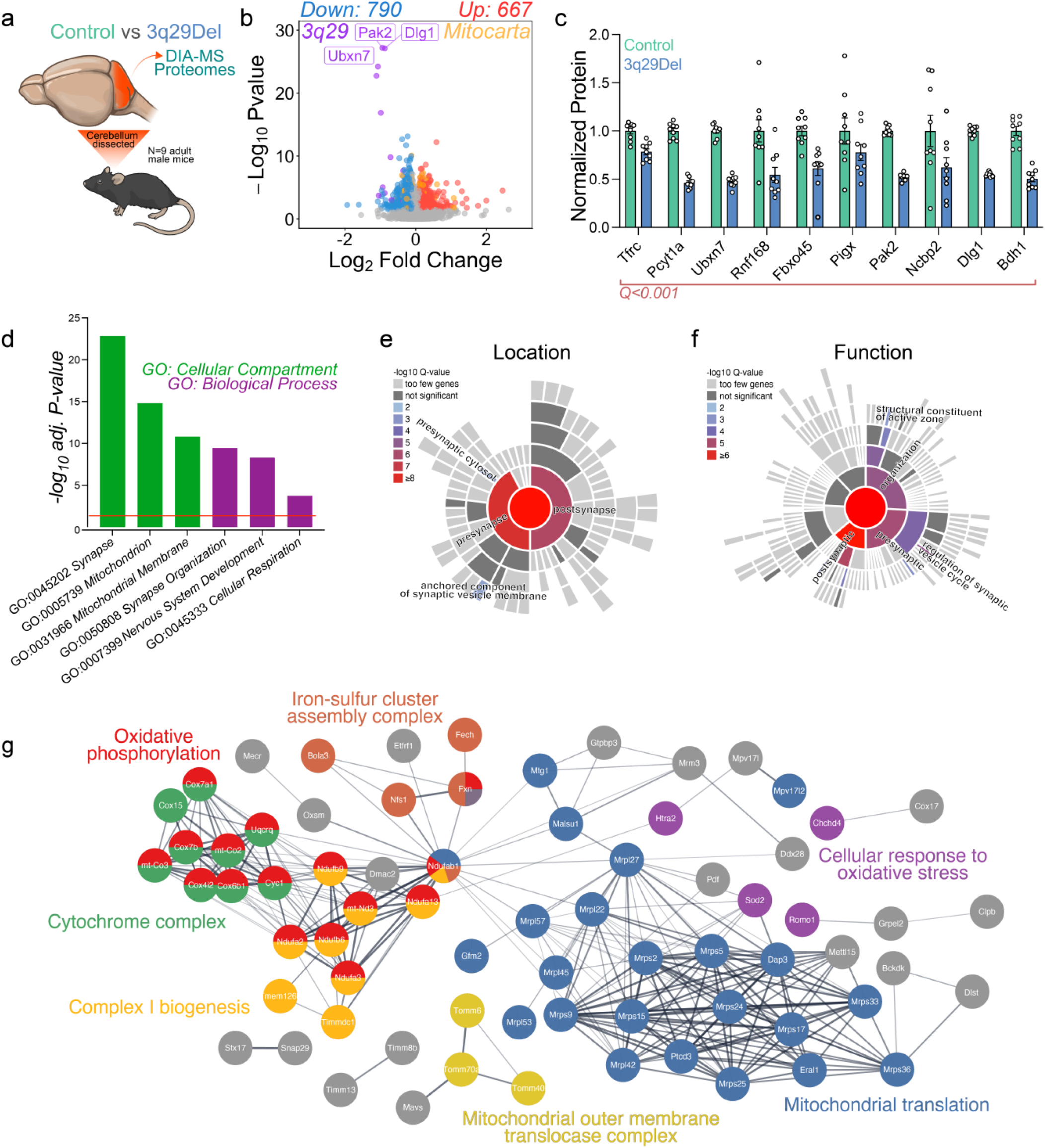
Proteomic and functional impacts of 3q29Del in cerebellum implicate disrupted synaptic and mitochondrial mechanisms. Experiment illustration (a) and volcano plot (b) of proteomic analysis of 3q29Del mouse cerebellum (N=9/genotype). 10 of 22 3q29Del interval proteins were detected (c) and all were significantly reduced in 3q29Del samples compared to WT controls. Summary of pathway analysis (d) of all proteins at q-value threshold <0.001. Evidence of synaptic enrichment prompted SynGO analysis of location 1. and function (f). Differentially abundant murine proteins (q<0.05, Abs.Log2FC>0.1) found in Mitocarta3.0 were searched in STRING v12.0 for physical interactions and functional enrichments linked to interacting nodes are listed.

To further understand specific aspects of mitochondrial protein dysregulation, we analyzed differentially abundant cerebellar proteins that are included in Mitocarta3.0^60^ with STRING v12^61^ for known physical interactions and functional enrichments (Fig. 4g). Remarkably, we found evidence for a highly connected network including many components of mitochondrial ribosomes, oxidative phosphorylation complexes I and IV, iron-sulfur cluster biogenesis factors, and cellular responders to oxidative stress.

To validate these findings and test for functional differences, we used high resolution respirometry in cerebellar mitochondria isolated from a new cohort of adult Control and 3q29Del mice (Fig. 5a-f). We found that oxygen flux (*J*O_2_, respiration) was significantly reduced under both non-phosphorylating (Fig. 5c) and maximum energy demand conditions (Fig. 5e). Furthermore, we found that electron leak (indicative of reactive oxygen species formation) was significantly increased in both states (Fig. 5d,f). As an integrative measure of oxygen consumption, electron leak gives a dynamic view into the fitness of 3q29Del mitochondria without bioenergetic demand (non-phosphorylating respiration) as well as at maximal levels of demand (ΔG_ATP_ -12.94). These outcomes are contrasted to traditional outcomes of oxidative stress generation that occur only at non-phosphorylating respiration, where ROS is artificially increased beyond physiological relevant levels due the absence of bioenergetic demand. These results indicate substantial perturbations of mitochondrial structure as well as the efficiency and integrity of the electron transport chain when bioenergetic demand is increased, which aligns with proteomic results. These data indicate that 3q29Del alters the proteome providing the basis for compromised mitochondrial respiration and increased oxidative stress in the mouse cerebellum.

**Figure 5.**
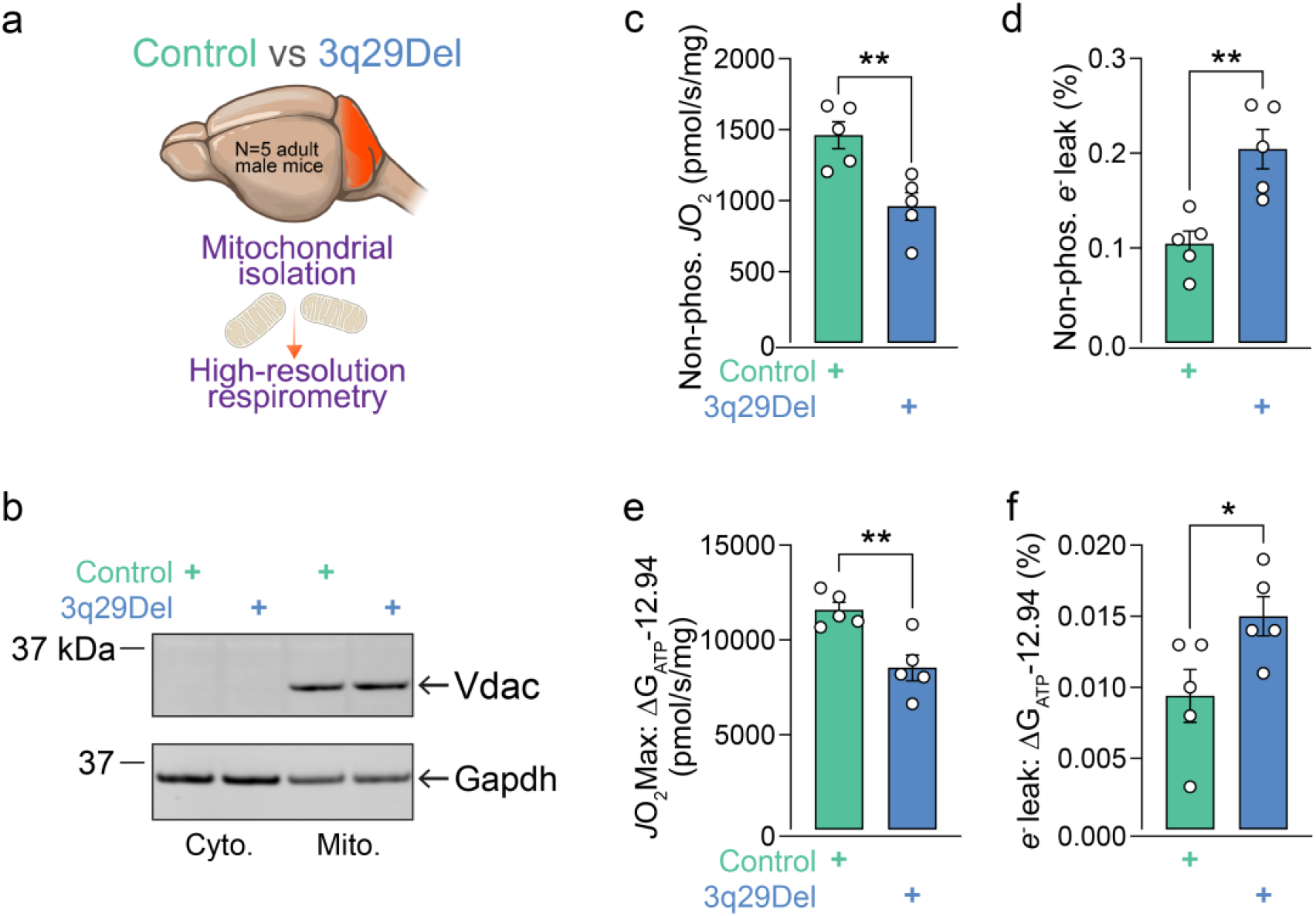
Mitochondria were isolated from N=5 WT and 3q29Del mice (a-b) for high resolution respirometry (c-f). Under both non-phosphorylating (c, p=0.0059; d, p=0.0038) and maximal respiration conditions (e, p=0.0047; f, p=0.0419), oxygen flux (*J*O^2^) was significantly reduced, and electron leak was significantly increased.

## 4. Discussion

In this study, we report novel cerebellar phenotypes in a mouse model of 3q29Del at structural, behavioral, and molecular levels. Using complementary methods in mice from two institutions, we found consistently reduced cerebellar size in 3q29Del mice, recapitulating the human phenotype of cerebellar volumetric reduction. Additionally, key cerebellum- and 3q29Del-associated behavioral phenotypes are conserved in mice, such as early vocal impairment, fine motor deficits, and decreased social communication. Finally, we found dysregulation of synaptic and mitochondrial proteins along with profound mitochondrial dysfunction in the 3q29Del mouse cerebellum. The observed changes in mitochondrial function build on previous observations in human neocortical organoids^8,9^, suggesting that this mitochondrial phenotype is a reproducible finding across species and brain regions. Together, these findings provide strong evidence that the cerebellum is impacted in 3q29Del mice, and that the cerebellum may be involved in 3q29Del pathology.

Mechanistic research to date has made use of transgenic animal models of ASD and SZ as well as human cellular and organoid experimental systems. Human cell cultures permit a fully human genetic background, and they are advantageous in elucidating cellular pathologies of genetic variants. However, they are limited in the size and developmental stage they can reach and cannot give much information on brain-wide circuits and structural phenotypes. Animal models allow the study of living mammalian brains with the caveat that some circuitry may not be fully conserved in humans. The relevance of transgenic animal models to complex neurological disorders has been questioned, but ultimately animal models are an important complement for basic and translational research^62^. In this study, we observed proteomic signatures indicative of mitochondrial phenotypes, which were validated in functional assays. We found dysfunctional mitochondrial respiration and reactive oxygen species buffering in the 3q29Del mouse cerebellum, which significantly builds on previous findings from multiple studies in 3q29Del animal and human cellular models^8,9,63^. Based on these results, this 3q29Del mitochondrial phenotype appears to be robust and conserved across species and cell types. Thus, the involved cellular pathology in the mouse model is more likely to be translatable to the human brain.

From a purely genetic lens, mitochondrial phenotypes would not necessarily be expected in 3q29Del, as only one gene on the 3q29 locus, *Bdh1*, encodes a mitochondria-localized protein^64^. Proteomic profiling revealed a broad disruption of proteins related to mitochondrial form and function in 3q29Del cerebella relative to control, consistent with the reduced respiratory capacity and increased electron leak observed in these mice. Filtering the differentially expressed proteins against Mitocarta further localized this phenotype to core mitochondrial processes, including electron transport chain (ETC) assembly, mitochondrial translation and replication, quality control, redox signaling, and maintenance of organelle integrity. Notably, proteins involved in complex I, cardiolipin, and cytochrome assembly and maintenance were dysregulated in 3q29Del mice, providing a potential molecular basis for impaired oxidative phosphorylation. Given the hemizygosity of *Tfrc*, which encodes the transferrin receptor 1, caused by 3q29Del, it is intriguing that the abundance of mitochondrial iron- and heme-regulatory proteins, including *Fech, Fxn, Isca1*, and *Tmem14c*, are altered. Together, these findings suggest that disruption of mitochondrial integrity and proteostatic maintenance may represent an upstream feature of 3q29Del that precedes, and potentially contributes to, impaired OXPHOS and redox homeostasis.

Notably, mitochondrial dysfunction is increasingly reported in 3q29Del as well as other ASD- and SZ-linked copy number variants such as 22q11.2 deletion (22qDel)^9-13^. Emerging evidence also indicates that these mitochondrial phenotypes are directly relevant to neuropsychiatric disease. In iPSC-derived neurons from individuals with 22qDel, mitochondrial dysfunction was found to be significantly worse in neurons from individuals with 22qDel and SZ compared to 22qDel and no history of SZ^11^. Furthermore, the rate of mitochondrial disease in individuals with ASD is substantially higher than the general population^6,7^. A direct connection from dysregulated cellular metabolism to elevated risk for ASD and SZ has not yet been established, but use of 3q29Del models to study this possibility will likely yield results that are applicable to other genetic variants associated with neurodevelopmental disability.

In addition to sharing cellular phenotypes with human cellular models, 3q29Del mice share key structural and behavioral phenotypes with the human 3q29Del population. Reduced cerebellar volume is a common and striking neuroanatomical difference in people with 3q29Del^14,15^, and we observed similar changes in 3q29Del mice. While the causes and functional consequences of reduced cerebellar size in 3q29Del are unknown, our results establish the possibility of studying these mechanisms in mice. We observed a reduction in cerebellar size that did not appear to vary by region. Human studies have also observed global size reduction in the cerebellar cortex^14,15^, but interestingly have reported a greater variability and magnitude of cerebellar volumetric reduction than we found in 3q29Del mice. Additionally, an anterior-to-posterior gradient in effect size has been observed in humans, with an apparently greater size reduction in more anterior cerebellar regions^14^, which we also did not observe in mice. These discrepancies may be due to polygenic and/or environmental interactions: much of cerebellar development occurs postnatally, making the cerebellum more vulnerable to early life environmental factors (such as injury/illness, stress, or malnutrition) than other brain regions. We controlled for mouse genetic background and many aspects of early life environment, likely reducing these causes of variability. The lower magnitude of the size difference we found suggests that cerebellar hypoplasia in people with 3q29Del may have an environmental component that was not present in our mice.

Some 3q29Del mouse behaviors also resemble human 3q29Del phenotypes. We investigated developing motor and vocal reflexes in 3q29Del mouse pups, and we found a reduction in call complexity that may be analogous to the speech delays commonly seen in children with 3q29Del^5^. Interestingly, we did not find a significant reduction in call number as was reported recently^50^. We also found no statistically significant difference in early motor reflexes in 3q29Del pups, but several animals in the 3q29Del group showed very long response times for the righting reflex, which was not observed in any wild-type control animals. This may suggest that 3q29Del alone is not sufficient to cause motor delays but does increase vulnerability to environmental factors.

Motor phenotypes in adult 3q29Del mice also strongly resemble human phenotypes. We observed no differences in gross motor coordination, which is consistent with what is known in 3q29Del population^15^. However, we did observe impairments in fine motor control. Many people with 3q29Del have writing disabilities^5^ and impaired visual-motor integration in tasks that require fine motor coordination^18^. We found that 3q29Del mice make more errors in a string pull test; specifically, they make more partial misses, which we found to be the most common error among all animals (Fig 4). Mice with disrupted cerebellar circuitry make more errors in string pull assays, but interestingly, those errors are more often full misses and double pulls^54^. Full misses and double pulls slow string pulling down more than partial misses (Supplemental video 1) and perhaps indicate greater motor difficulty. This result indicates that 3q29Del produces a more mild, but still significant, impairment than the manipulations used in previous studies.

This study has several limitations. First, while structural and functional differences in the cerebellum were observed, 3q29Del is a global manipulation, not cerebellum specific. Thus, it is possible that dysfunction in other brain regions contributes to the observed behavioral changes. We demonstrated that the mouse equivalent of a global 3q29 deletion (as is present in the human 3q29Del syndrome) recapitulates key human 3q29Del phenotypes, but the direct mechanistic causes of these phenotypes still must be investigated. Second, we were unable to test adult vocalizations beyond a courtship assay, which involves only male mice and is primarily a social setting. Inclination to vocalize (number of calls) can be interpreted as sociability, but we do not know whether the reduced call complexity we observed also reflects reduced social interest, or whether the 3q29Del adults have a vocal impairment. Finally, while we selected multiple time points associated with developmental milestones of interest, we did not conduct detailed experiments across the full mouse developmental timeline within the scope of this study. Thus, there may be gaps in which the effects of 3q29Del are more prominent.

Here we have characterized translationally relevant cerebellar phenotypes in a mouse model of 3q29Del. We have also highlighted mitochondrial dysfunction as an avenue of further mechanistic study: the 3q29Del mitochondrial phenotype is not only conserved across species, but is also similar to mitochondrial outcomes in other ASD- and SZ-linked copy number variants, potentially implicating mitochondrial dysregulation in the cellular pathology of these disorders. Additionally, studying the cerebellum provides a strong foundation for understanding the impact of neurodevelopmental disorders on the entire brain. Cerebellar circuitry is highly conserved between mice and humans^47,65^, and many cerebellar phenotypes of 3q29Del are recapitulated in the mouse model. Thus, the cerebellum is a promising region for research to link biological processes, such as mitochondrial function and structural development, to behaviors. Understanding these processes and mechanisms may lead to interventions that can mitigate symptom severity and improve quality of life for those born at risk.

## DISCLOSURES

### Declaration of generative AI and AI-assisted technologies in the manuscript preparation process

During the preparation of this work, the authors used Copilot and ChatGPT to troubleshoot code used to organize proteomic and vocalization data. The authors reviewed and edited the code as needed after using these tools and take full responsibility for the content of the published article.

